# Biological research in Papua New Guinea: a publication activity analysis

**DOI:** 10.1101/2021.08.25.457669

**Authors:** Vojtech Novotny, Pagi Toko

## Abstract

The Nature Index and the Web of Science databases are used to analyse the publication patterns in life sciences in Papua New Guinea, aiming to identify the ways of improving the country’s research output.

## Introduction

New Guinea is the floristically richest island in the world, harbouring approximately 5% of global plant and animal diversity, as well as the third largest rainforest in the world (Novotny & Molem 2020. Nature 584:531). This diversity is also matched with the diversity of human pathogens and parasites, making the island an attractive area for biodiversity and medical research. However, its diversity remains poorly known. In Papua New Guinea (PNG), occupying the eastern part of the island, the indigenous knowledge of the natural world is being rapidly lost (Kik et al. 2021. PNAS 118, e2100096118) while the intensity of research is not adequate to the magnitude of the country’s biodiversity or its conservation importance.

PNG had started with two universities at the time of its independence in 1975, the University of PNG and the PNG University of Technology. The list of its universities has grown to eight at present. They cater to a rapidly increasing population, from 3.1M in 1975 to 9.0M at present. The proportion of population with tertiary education is increasing only slowly, presently at 6.6% of the 30-34 years old cohort with tertiary education (PNG National Statistical Office, 2019 Demographic and Health Survey).

The country has also a network of applied research institutes. In biological sciences, it includes the PNG Institute of Medical Research (PNG MRI), PNG Forestry Research Institute (PNG FRI), PNG National Agriculture Research Institute (PNG NARI), as well as the PNG Coconut and Cocoa Research Institute and PNG Coffee Research Institute, specializing on cash crops. The PNG National Research Institute contributes to social and political aspects of biodiversity use and conservation research. This structure was established around the time of the country’s independence and does not reflect later developments, such as the increased importance of biodiversity and conservation research or new cash crops, such as vanilla.

The museums are represented by the PNG National Museum and Art Gallery. The NGO scene in PNG is focused primarily on practical improvements of livelihoods, health, education and conservation, with research sometimes being a smaller part of their focus. The New Guinea Binatang Research Center (BRC) is an exception from this rule, focusing primarily on biodiversity research.

The present analysis is an attempt to map the current research productivity in PNG as a basis for the assessment of the ways to improve it in the future.

## Methods

This analysis is based on the publications included in the Nature Index (https://www.natureindex.com/) and the Web of Science (WoS) (https://www.webofscience.com/) databases. The Nature Index focuses on the highest quality papers while WoS is more comprehensive. While the Nature Index is publicly available, the access to WoS requires subscription. Very few, if any, PNG institutions have access to WoS, illustrating already one of the problems of research in PNG.

The Nature Index was analysed for all life science publications, including medicine. Only a few papers from PNG a year qualify for the Index so that year-to-year results could be erratic. We therefore present only summary analysis for the entire period for which the index is available, 2015-2020 data. One measure of productivity is the share of the authors from a particular institution in the overall authorship (e.g., each author has 0.1 share on a paper with 10 authors), another is the number of papers included in the Nature Index irrespective of the authorship share.

The Core Collection of the WoS database was used to assess the institutional contributions to research productivity in PNG. It was quantified using the WoS papers with “Papua New Guinea” in the “topic” field, classified into the following subjects: zoology, plant sciences, environmental studies, entomology, ecology, environmental sciences, biodiversity and conservation, evolutionary biology, forestry, agronomy, fisheries, biology, ornithology, mycology and freshwater biology. These represent non-medical biological fields. The subset of the papers with at least one PNG author were then listed according to the participating PNG institution (note that one publication can be listed multiple times, depending on the number of PNG institutions involved). The analysis was performed for 2001-2020.

The share of local authors in non-medical biological publications was quantified as the proportion of papers with “Papua New Guinea” in the “topic” field with at least one author from a PNG institution. The same analysis was repeated for papers about Australia, and the share of their local authors.

The research intensity standardized by biodiversity present in the country was quantified as the number of WoS papers per bird species in each country, over the 1975 – 2020 period (i.e., the duration of the independent state of PNG). The papers were filtered using “bird” and the name of the country in the “topic” field. Only selected countries, representing a mix of small and large, species poor (Temperate) and species rich (tropical) ones were used.

The intensity of research of a particular ecological phenomenon in PNG and elsewhere was examined using long elevational gradients in the tropics, comparing the number of publications for Mt. Kinabalu (Malaysia), Mt. Kilimanjaro (Tanzania), Mt. Cameroon (Cameroon) and Mt. Wilhelm (PNG). These are all tropical mountains reaching the alpine zone, also well known as the highest peaks of their respective countries. The 2001-2017 papers, and their citations, with the name of the mountain included in the “topic” field were considered, separated into those on biodiversity topics and other subjects.

## Results

The performance of PNG institutions during 2015-2020 shows an overwhelming dominance of the PNG IMR that is responsible for almost 70% of the PNG productivity measured in authorship share (Fig. 1). The PNG MRI is followed by BRC, and further by the PNG National Museum and the University of PNG. Additional six institutions that produced at least one Nature Index paper in the examined period have only negligible authorship shares. The top two institutions, PNG MRI and BRC, retain their leading position also when the number of papers is considered, followed by the University of PNG in the third place (Fig. 1).

**Fig. 1.**
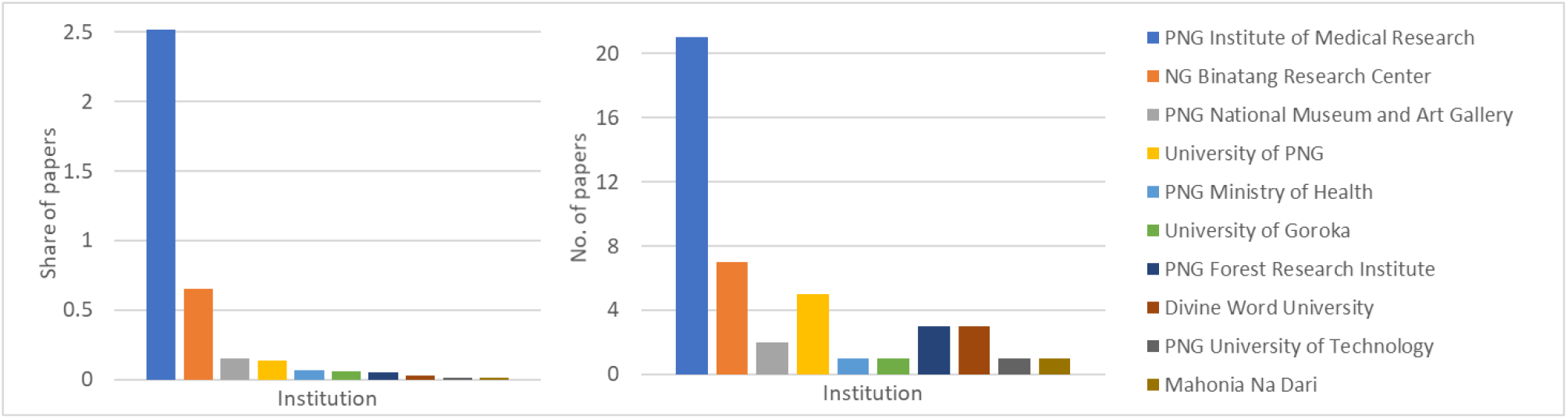
High quality publications (included in the Nature Index) in life sciences produced by PNG institutions in 2015-2020; the authorship share (L) and number of publications (R) are given for all institutions included in the Index.

PNG research in sciences is entirely dominated by life sciences, representing 45 out of 46 papers included in the Nature Index, with a paper from the Rabaul Volcanic Observatory representing the only additional contribution for the Earth Sciences / Chemistry / Physics fields. Our analysis thus also reflects the overall PNG research productivity beyond life sciences, again dominated by the PNG IMR, followed by BRC.

The top seven institutions publishing in non-medical biology in PNG produced 238 publications in 2001-2000 according to the more comprehensive WoS database. Overall, the University of PNG published the highest number of papers (64), followed by the BRC (57) and, more distantly, the PNG NARI (33) (Fig. 2). The top seven institutions include two universities, two research institutes, two NGOs and one government agency. The trends in the last five years compared with the previous 15 years show marked decline of the publication leader, the University of PNG, and a rapid increase of BRC that has become the leading institution during the 2016-2020 period.

**Fig. 2.**
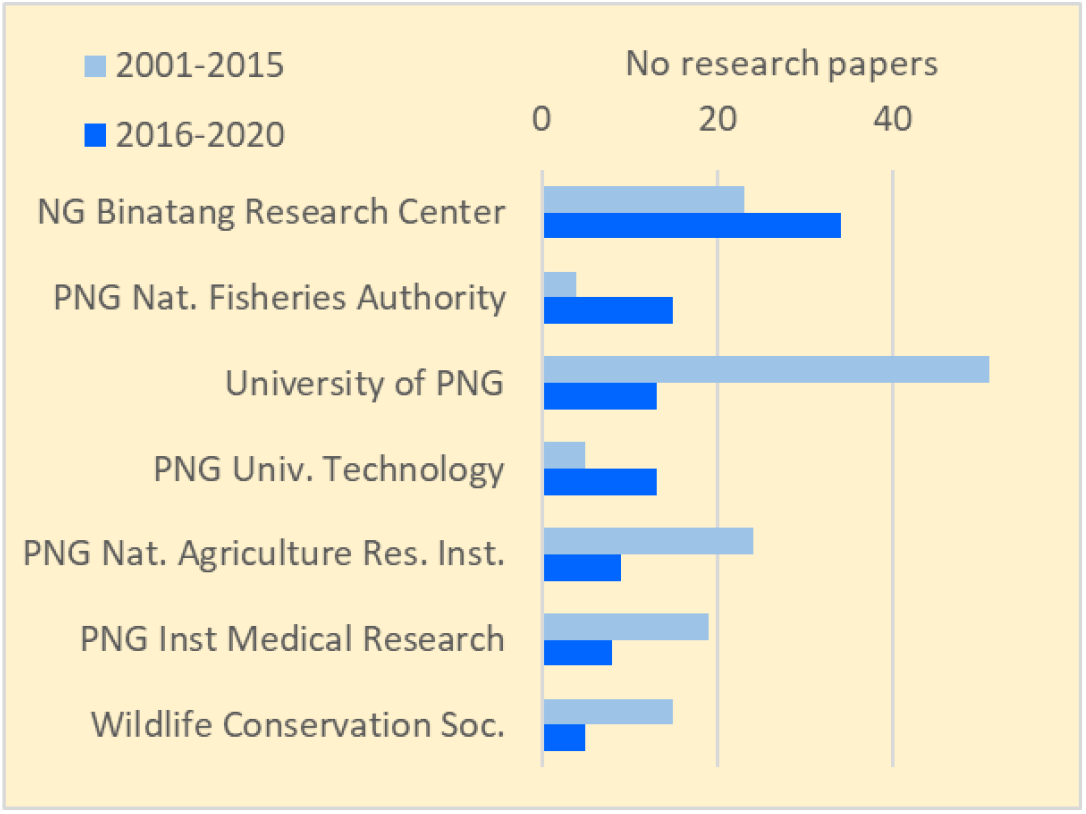
Number of non-medical biology publications on Papua New Guinea published by PNG institutions in the past 20 years (publications included in the WoS database).

The publication output of papers on PNG in non-medicinal biology has been 18x smaller than that on Australia during the past 20 years (Fig. 3). Only 14% of papers on PNG had at least one local author (with a PNG address), while for Australia the share of at least partially locally-authored papers was 76%. This means that Australian researchers participated on 93x more papers on the biology of their own country than PNG researchers did. There appears to be a slow increase in the share of local authorship over the years, both in PNG and Australia (Fig. 4).

**Fig. 3.**
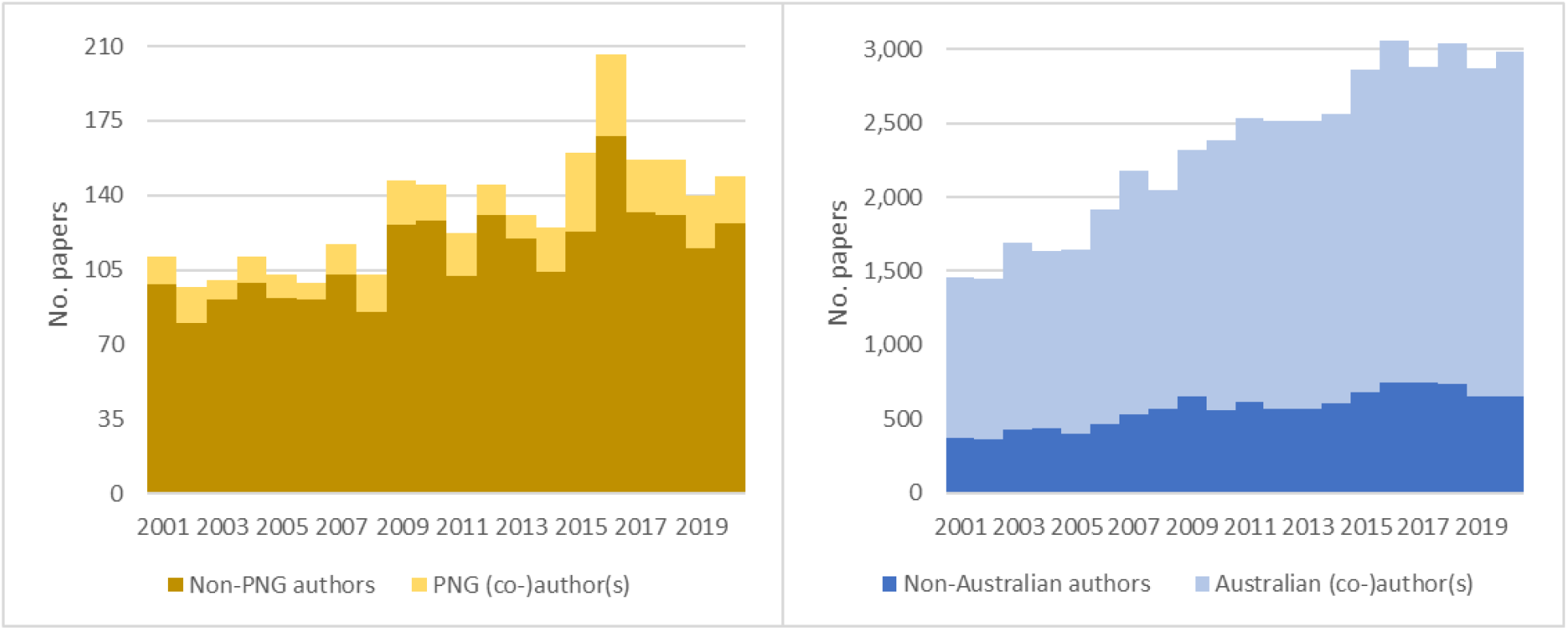
Number of non-medical biology publications in WoS database on the topic of Papua New Guinea (L) and Australia (R) that are (co-)authored by local researchers or published exclusively by foreigners.

**Fig. 4.**
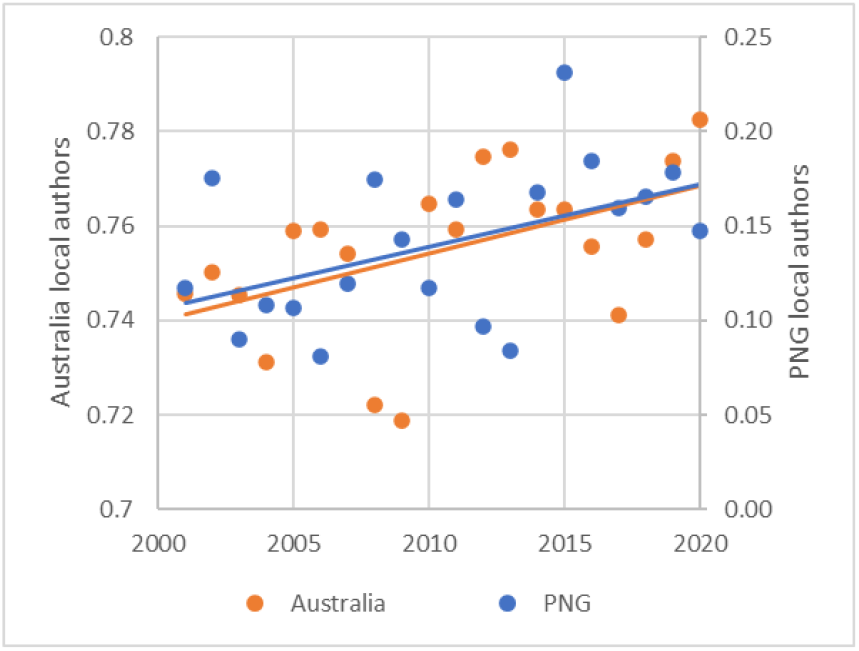
The proportion of papers with at least one local (co-)author published annually about PNG and Australia.

Australia is leading, among 20 arbitrarily selected countries, in the number of ornithological publications per local bird species in 1975-2020 (Fig. 5). Its publication rate of >5 papers per species is higher than in species-poor countries, such as Germany or Sweden. Most of species-rich tropical countries, including PNG, reached an order-of-magnitude lower publication rates than Australia.

**Fig. 5.**
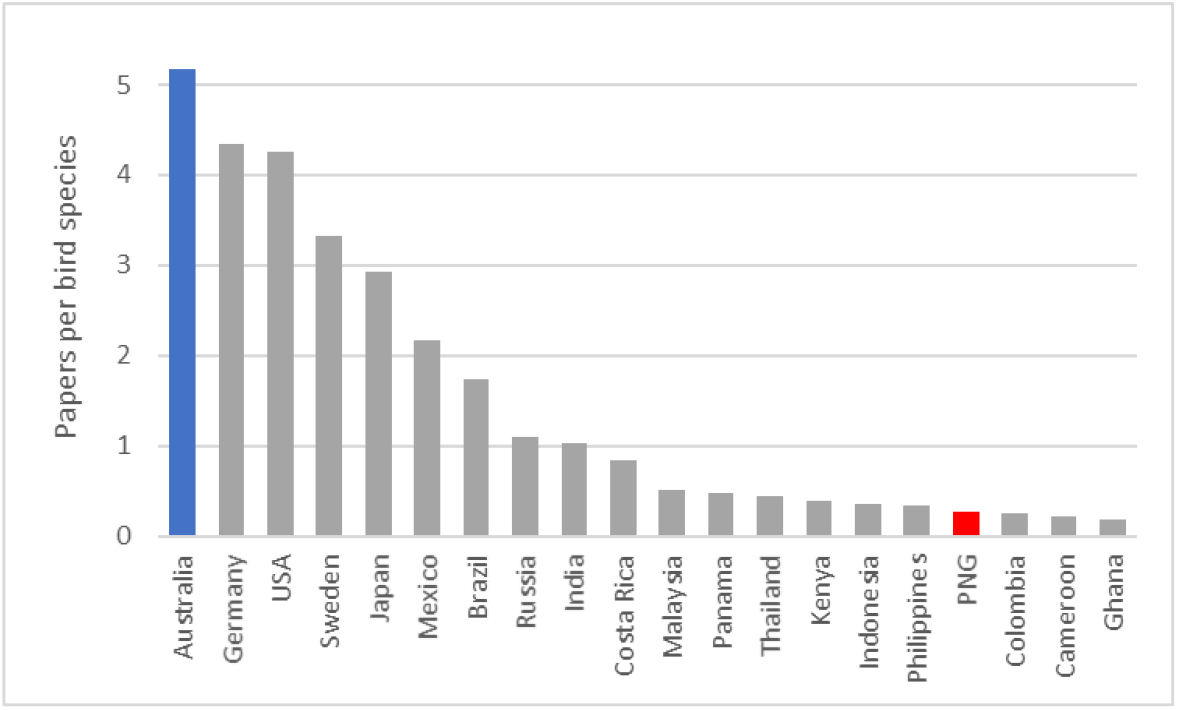
The number of ornithological papers per local bird species published in a set of 20 countries in 1975-2020.

The number of publications and citations overall, and on biodiversity, was markedly lower for Mt. Wilhelm than for any of the other three prominent elevation gradients in the Palaeotropics (Fig. 6).

**Fig. 6.**
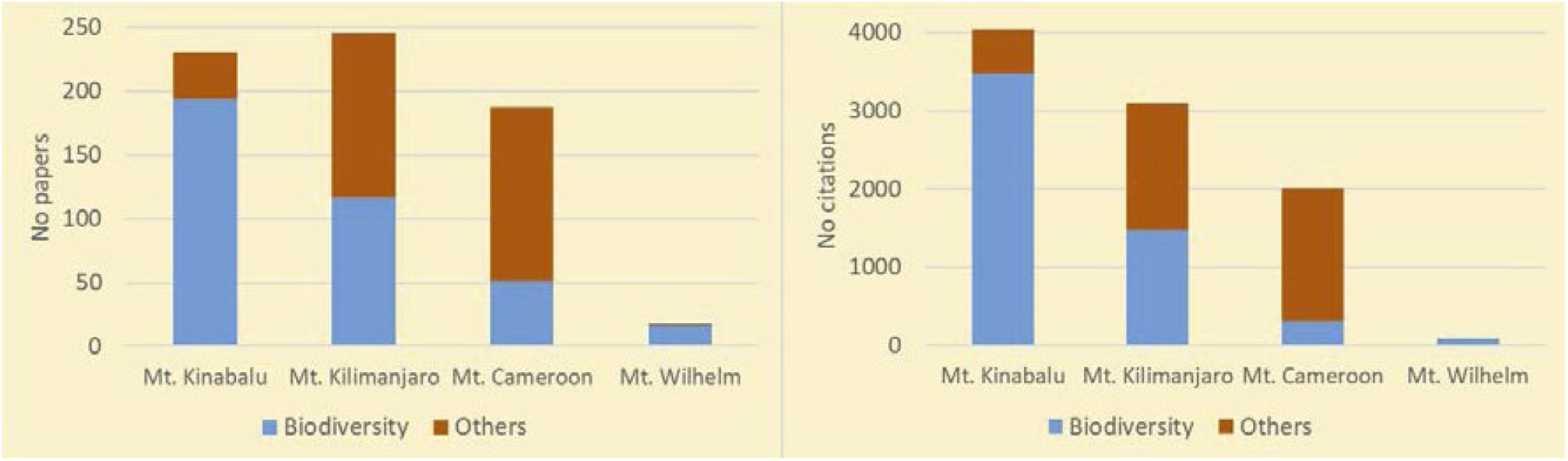
The number of WoS publications (L) and citations (R) on the four elevation gradients reaching alpine zone in the tropics.

## Discussion

The results of this analysis are partly predictable, partly surprising. On the positive note, the analysis demonstrate that it is possible to create and maintain a highly productive research institution in PNG, as exemplified by the PNG Medical Research Institute. Further, it is clear that at least in biodiversity research, an NGO may have a significant impact on the national research productivity, as demonstrated by the NG Binatang Research Center.

Less optimistically, the analysis demonstrates the weakness of PNG universities. Even the relatively high productivity of the nation’s premier University of PNG is mostly the thing of the past, as its performance in the last five years, measured by both the Nature Index and WoS, clearly shows. The other universities have yet to make their name in biological research. The performance of the PNG research institutes other than PNG IMR should be more prominent as well, considering that their staff should be fully occupied by research.

There are several underlying problems determining the low performance of PNG science. The local universities are barely keeping pace with increasing population (the median age in PNG population is 22.4 years) and are essentially teaching universities where the low staff-to-students ratio demands most of the staff time to be spent on teaching. The postgraduate programmes are limited and expensive, with no or very few studentships available. This leads to very low number of MSc and PhD graduates, in biology typically less than 10 MSc degree awarded annually for the entire country. However, postgraduate students are the main workforce of science, so the lack of the critical numbers of postgraduate students means there is nobody at the universities to work on more ambitious research projects.

The situation should be better at research institutes. However, their productivity is harmed by their almost complete isolation from university students, who could bring new ideas and enthusiasm. Another serious problem affecting all institutions in PNG is outdated system of research funding that allocates resources to institutions (universities and research institutes) by administrative decision, rather than to freely competing research projects via a national grant agency. The current funding system does not reward productivity.

Unsurprisingly, the intensity of biology research in PNG is lower than in the neighbouring Australia. Australia has 2.5x higher population, 56x higher national GDP, 33x higher when adjusted to the purchasing power parity (PPP). The 93x lower performance of PNG than Australian researchers means that PNG science under-performs Australia 2–3x, depending on whether we consider GDP or the PPP-adjusted GDP to be the best predictor of research performance. This is the lag that could conceivably be closed, with a focused effort. A higher participation of PNG researchers in the research of their own biodiversity would close this gap.

The higher participation cannot and should not be achieved by administrative measures, such as by mandating all overseas scientist to include PNG colleagues as co-authors as some other tropical countries, including Indonesia, sometimes do. The overseas researchers should be strongly encouraged to involve their local counterparts, while there must be also more extensive and intense training for science in PNG so that such collaborations could be genuine and beneficial to both sides. The goal is certainly not exclusive publication by Papua New Guineans on PNG biology. Rather paradoxically, extended collaborations rather than isolation will lead to improved performance by local scientists, as the highly collaborative programme of the PNG Institute for Medical Research shows.

There is a large unexploited potential for research of PNG biota. For instance, PNG ornithological research would have to increase 20x before reaching the same intensity, in terms of publications produced per bird species, as in Australia. Likewise, elevation gradients, eminently important as natural laboratories for the study of climate change on biota, remain virtually unexplored in PNG in comparison with other tropical regions. This is despite the fact that Mt. Wilhelm, together with the neighbouring Saruwaged Range, qualifies one of the seven floristically richest areas in the world (Barthlott et al. 2007, Erdkunde 10: 305).

Hosting and collaborating with overseas research projects could not only improve local expertise in PNG but also become a profitable industry, bringing employment and “research tourism” (West 2008, Current Anthropology 49: 597). While PNG has so far failed to develop as a tourist destination (it is a destination for only 0.01% of the world’s tourists; www.theglobaleconomy.com), it has an excellent potential to be an internationally competitive user-friendly destination for international research. The PNG system of research visa and permits is set up and working better than in most other tropical countries, and widespread use of English is also advantage. PNG is thus a competitive destination for international research, but so far seriously under-utilized except for medical sciences.

## Conclusions

PNG research productivity in life sciences is good in the medical research due to the PNG MRI, but unsatisfactory in the other areas. PNG universities need to develop a more accessible postgraduate training programs to create a workforce for more ambitious research projects. The country’s structure of research institutes needs an update, including biodiversity and conservation research, as well as new cash crops. The research institutes need to open to students. A national grant agency, funding research projects on competitive basis, is needed in addition to the current direct funding of institutions. PNG has a great potential to develop as an internationally competitive destination for life science research which could bring both expertise and revenue to the country.

